# Myosin-II activity generates a dynamic steady state with continuous actin turnover in a minimal actin cortex

**DOI:** 10.1101/312512

**Authors:** Sonal, Kristina A. Ganzinger, Sven K. Vogel, Jonas Mücksch, Philipp Blumhardt, Petra Schwille

## Abstract

Dynamic reorganization of the actomyosin cytoskeleton allows a fine-tuning of cell shape that is vital to many cellular functions. It is well established that myosin-II motors generate the forces required for remodeling the cell surface by imparting contractility to actin networks. An additional, less understood, role of myosin-II in cytoskeletal dynamics is believed to be in the regulation of actin turnover; it has been proposed that myosin activity increases actin turnover in various cellular contexts, presumably by contributing to disassembly. *In vitro* reconstitution of actomyosin networks has confirmed the role of myosin in actin network disassembly, but factors such as diffusional constraints and the use of stabilized filaments have thus far limited the observation of myosin-assisted actin turnover in these networks. Here, we present the reconstitution of a minimal dynamic actin cortex where actin polymerization is catalyzed on the membrane in the presence of myosin-II activity. We demonstrate that myosin activity leads to disassembly and redistribution in this simplified cortex. Consequently, a new dynamic steady state emerges in which actin filaments undergo constant turnover. Our findings suggest a multi-faceted role of myosin-II in fast remodeling of the eukaryotic actin cortex.

## INTRODUCTION

Rapid and controlled modulation of cell shape provides the foundation for various cellular functions, such as division, growth, migration and adhesion. In eukaryotic cells, much of this modulation can be attributed to the dynamics of the actomyosin cortex that lines the inner surface of the plasma membrane. The cortex is a thin network of actin filaments associated with myosin motors (see Salbreux et al., 2012 for a comprehensive review). It is further reinforced by a collection of actin binding proteins that mediate filament-filament and filament-membrane interactions. Additionally, myosin-II motors drive contractility of the actin meshwork, generating forces at different length scales that enable fast remodeling of the cell surface.

Another factor that imparts dynamicity to the cortex is the continuous turnover of all the protein components of this network. Relative rates of turnover can determine the mechanical properties of the cortex and are, therefore, subject to local regulation. For instance, actin turnover has significant consequences on cortical tension (Tinevez et al., 2009) and various network elements are implicated in imparting different turnover rates to subpopulations of actin (Fritzsche et al., 2013). The rate of actin turnover even within the same cell has been shown to vary between functionally distinct regions of the cortex (Murthy and Wadsworth, 2005). Turnover rates of actin can be tuned by manipulation of either assembly or disassembly processes. Lipids in the bilayer can orchestrate local polymerization via activation of nucleators such as the Arp2/3 complex (Higgs and Pollard, 2000). Disassembly, on the other hand, is believed to be enhanced by myosin activity itself, although dedicated severing proteins also exist (Blanchoin et al., 2000). Suppression of myosin activity was implicated in reduced actin turnover in a range of cellular contexts (Guha et al., 2005, Medeiros et al., 2006, Murthy and Wadsworth, 2005, Yang et al., 2012).

Due to the complexity of cellular environments, *in vitro* experiments have proved more amenable for direct observation of myosin-II driven severing of actin filaments (Haviv et al., 2008, Murrell and Gardel, 2012, Vogel et al., 2013) as well as network disassembly (Reymann et al., 2012). While various other reconstitution studies have addressed the mesoscopic effects of myosin contractility on actin network reorganization, a direct investigation of the role of myosin in network turnover has been lacking. The observation of filament turnover has previously been limited either by factors like stabilized actin filaments, diffusion constraints, restricted polymerization (Koster et al., 2016, Linsmeier et al., 2016, Murrell and Gardel, 2012, Smith et al., 2007, Stam et al., 2017, Vogel et al., 2017, Vogel et al., 2013); or by a focus on network-level changes rather than individual network components (Carvalho et al., 2013, Soares e Silva et al., 2011, Kohler et al., 2011, Bussonnier et al., 2014).

By using a biomimetic actin cortex with minimal restraints on the mobility of components and coexistence of assembly and disassembly, we demonstrate that myosin-II activity can generate turnover in membrane-associated dendritic actin networks. This observation was enabled by the following features incorporated during reconstitution: 1) polymerization driven by catalysts localized to the membrane and 2) no additional anchoring of either the myosin or the actin. While sharing qualitative attributes of the contraction process with previous studies, our minimal cortex additionally showed breakdown of actin networks on the membrane, redistribution of network components and finally, recruitment of the released components into new assembly--altogether, the essential aspects of filament turnover.

## RESULTS

### Assembly of a dynamic actomyosin cortex on supported lipid bilayers

For the reconstitution of a biomimetic minimal cortex with the potential for actin network turnover, we required a strong catalyst of actin polymerization to oppose myosin-driven disassembly. The constitutively active version of the murine N-Wasp, also known as ″VCA″, served as the catalyst. This protein was anchored to a supported lipid bilayer (SLB) with the stable association of a 10x Histidine tag to nickelated lipids (Nye and Groves, 2008). Not only does the membrane localization of the catalyst emulate the cellular scenario, but it also allows us to preferentially image the polymerization process, by using Total Internal Reflection Fluorescence (TIRF) microscopy. VCA has been extensively studied *in vitro* and has been shown to promote the autocatalytic growth of a branched actin network, when combined with the Arp2/3 complex (Pantaloni et al., 2000, Rohatgi et al., 1999). Such ″dendritic″ network assembly is fast and, therefore, a suitable choice for counterbalancing disassembly by myosin activity. Muscle myosin-II motors were preassembled into filaments and introduced into the system after actin assembly (Fig. 1A).

**Figure 1.**
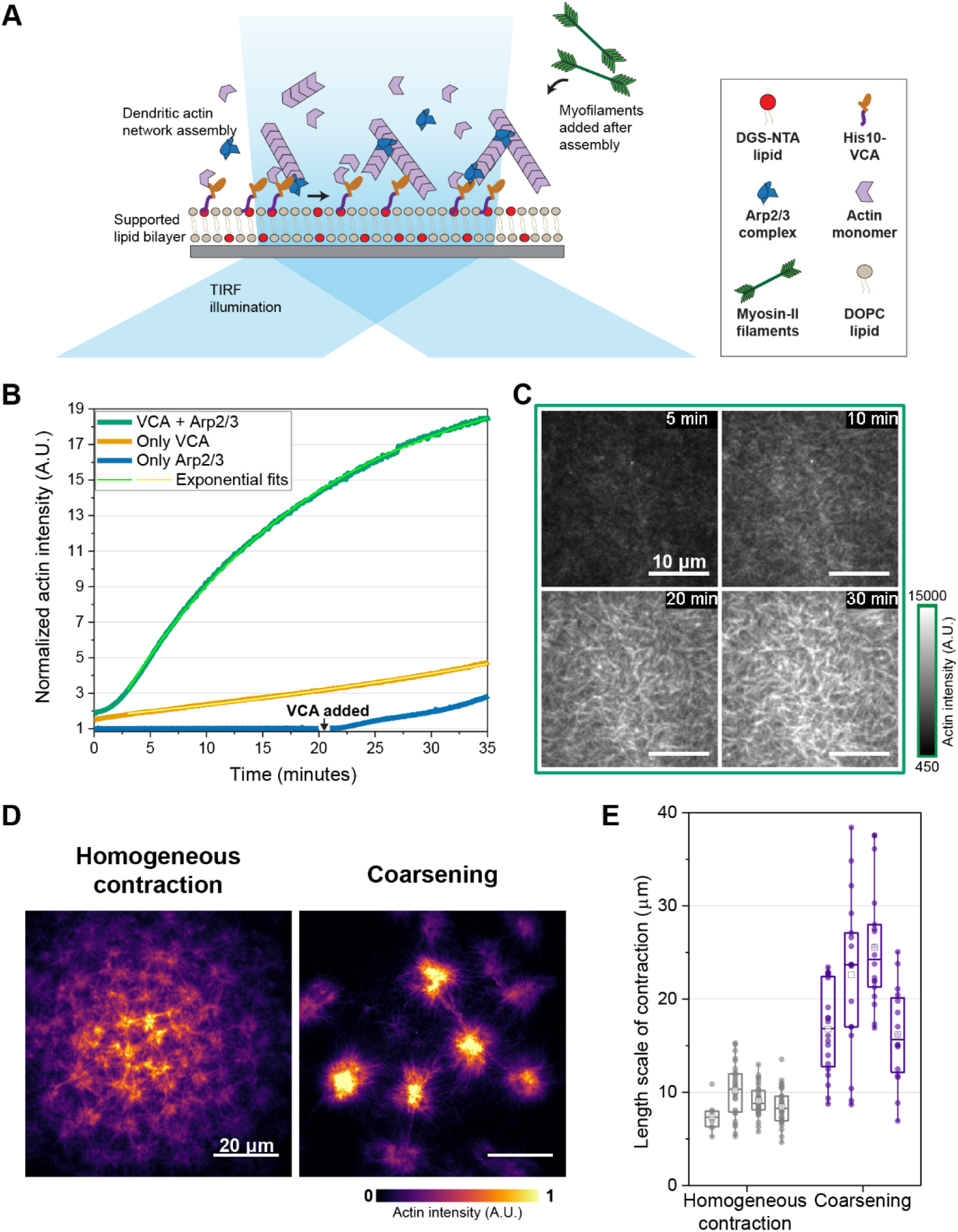
Assembly of a dynamic actomyosin cortex on supported lipid bilayers. A) Schematic depiction of the design of a minimal dynamic actomyosin cortex. A dendritic actin network is assembled on a supported lipid bilayer by Arp2/3-induced branch nucleation, catalyzed by VCA anchored to nickelated lipids (DGS-NTA). Myofilaments are introduced after actin assembly and the system is observed by TIRF microscopy. B) Time profiles of actin assembly on the membrane and its dependence on different reaction components. The growth is slower in the absence of Arp2/3 and undetectable in the absence of VCA. Addition of VCA (arrow) to the latter shows immediate increase in actin intensity. Concentrations used were 1 μM actin, 10 nM Arp2/3 complex and 100 μM ATP on an SLB with 1 mol% DGS-NTA. Fit parameters are provided in Table S1. C) Time frames from the profile of the VCA + Arp2/3 reaction depicted in B. Bundling can be observed at 30 minutes. Intensity scale is provided as reference for comparison with images of control conditions in Figure S1. D) Representative images of the two modes of actomyosin contraction observed in the presence of 0.8 μM myosin-II. The modes differ in density and distribution of condensates. E) Characteristic length scale of contraction calculated as nearest neighbor distance between foci. Plot shows 25-75 percentile as box, range within 1.5 interquartile region as whiskers, median as line and mean as square. Individual data points are depicted as closed circles. Length scales are significantly different between the two modes (p<0.05; Mann-Whitney test).

We started with the characterization of the actin assembly process in the absence of myosin motors. As expected, after a short time lag, surface actin intensity increased fast on the VCA-functionalized SLBs in the presence of Arp2/3 and actin monomers (Fig. 1B, Movie 1). The observed time profiles are characteristic of the autocatalytic nature of dendritic actin network assembly (Pantaloni et al., 2000). Actin network growth was dense and resulted in bundle-like organization of filaments with time (Fig. 1C). In comparison, actin assembly was significantly weaker when Arp2/3 was excluded from the reaction and nearly non-existent in the absence of VCA (Fig. 1B). While VCA alone was sufficient to induce some actin assembly, the resulting meshwork was coarser than in the presence of Arp2/3 (Fig. S1A). This Arp2/3-independent assembly can be attributed to the ability of VCA to enhance actin filament elongation on surfaces by increasing the local availability of monomers (Bieling et al., 2018). In contrast, Arp2/3 alone failed to initiate any actin polymerization on the membrane (Fig. 1B, Fig. S1A), since VCA not only activates the Arp2/3 complex, but is also necessary to mediate actin-membrane interactions in this minimal cortex. Consequently, adding VCA to this sample immediately leads to increased actin intensity on the surface and initiation of filament assembly (Fig. S1A). Altogether, these trends confirm that the actin assembly we observe on the membrane is VCA-dependent, with considerable contribution from Arp2/3-induced branching.

Interestingly, the diffusional mobility of the VCA on the membrane appeared to be essential for the observed actin network growth, as holes in the lipid bilayer were largely devoid of actin, even though VCA accumulated there (Fig. S2B). We also confirmed that VCA intensity on the membrane remained stable during actin assembly (Fig. S1C, Movie 1). Actin was grown for longer periods to bring the growth process close to saturation (Fig. S1D) before adding myofilaments.

Introducing myosin-II to the dendritic actin network resulted in condensation of actin into contraction foci. Interestingly, two distinct length scales of actomyosin contraction were observed under our experimental conditions (Fig. 1D, Movie 2). In some samples, the contraction foci were distributed homogeneously with shorter distances between them (9.1±2.2 μm) and actin enrichment in these foci was less pronounced. In other cases, we observed a distinct coarsening process with actin condensing into dense foci, separated by larger length scales (20.3±7.4 μm). The characteristic length scales of the two contraction modes were calculated as nearest neighbor distance of the contraction foci (Fig. 1E). The observation of multistage coarsening is in line with previous studies on disordered actin networks, both membrane-associated and in solution (Backouche et al., 2006, Kohler et al., 2011, Linsmeier et al., 2016, Murrell and Gardel, 2012, Smith et al., 2007, Soares e Silva et al., 2011, Stam et al., 2017, Vogel et al., 2013).

The bimodal outcome of contraction led us to investigate the factors that impact length scale of contraction in membrane-associated dendritic actin networks. We found that this length scale responded to manipulations in surface density of VCA, Arp2/3 concentration, myosin concentration and branch capping (Fig. S2). A five-fold increase in VCA surface density, leading to a net increase in membrane attachment of the actin network, reduced the length scale by one-third (Fig. S2A). Further, enhanced branching resulting from increasing Arp2/3 concentrations trended towards lower length scale (Fig. S2B). Lower myosin concentrations and the introduction of branch capping, presumably resulting in shorter filament lengths, also favored lower length scales (Fig. S2C-D). These dependencies corroborate a previous *in vitro* study, where membrane adhesion and filament cross-linking were shown to affect contraction length scale in a disordered actin network (Murrell and Gardel, 2012).

### Myosin-II breaks down membrane-associated actin networks

Irrespective of the length scale of contraction, we found that myosin activity led to disassembly of the actin cortex (Fig. 2). The initial stages of myosin contraction showed bundling of actin filaments, followed by a compaction of the actin network (Fig. 2A a-white arrows, Movie 3). As the actin condensed into contraction foci, rips in the actin network could be observed. Finally, this process resulted in a decrease in actin intensity on the surface, indicating network breakdown. Myofilaments were associated with the bundles as well as the condensed actin foci (Fig. 2A b, yellow arrows). They, however, accumulated on the membrane even as the actin intensity dropped with disassembly (Fig. 2B).

**Figure 2.**
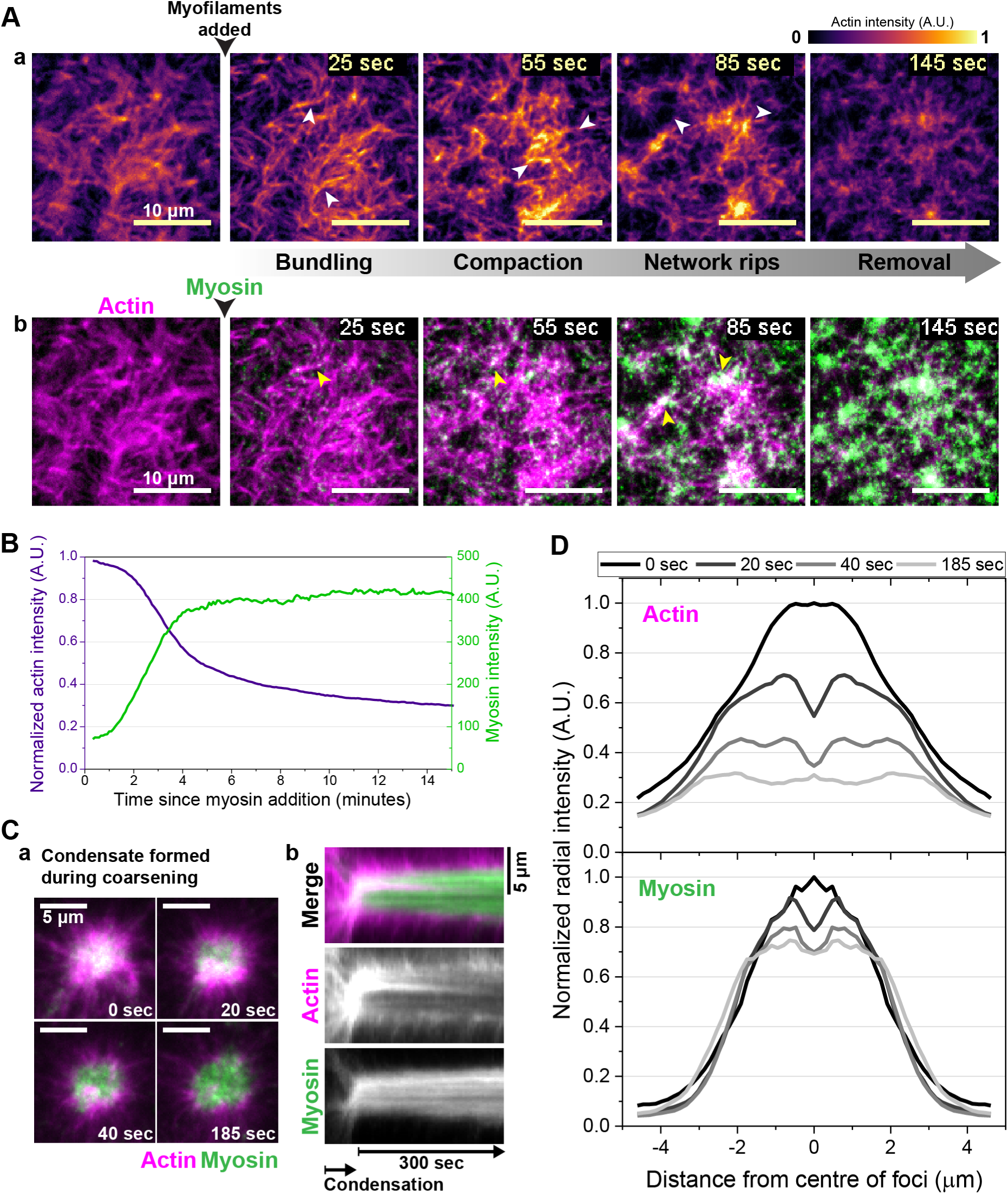
Myosin-II breaks down membrane-associated actin networks. A) After addition of myofilaments, dendritic actin networks show bundling, compaction, network rips and finally removal from the membrane (a). White arrows highlight the mentioned feature in each frame. Panel b shows the location of myofilaments (green) in the corresponding frames of panel a. Yellow arrows highlight myosin in actin bundles and foci. B) Time profiles of actin and myosin intensities from the sample described in A. The actin intensity is normalized to the value before addition of myosin and decreases as the myosin intensity on the surface increases. C) Selected time frames from TIRF imaging (a) and kymograph (b) depicting the release of actin from a single condensate formed during coarsening. The kymograph shows intensities averaged over a 10-pixel thick vertical line through the center of the condensate. Data is representative of 10 condensates observed with labeled myosin. D) Radially integrated intensities of actin and myosin at the different time points depicted in C showing hollowing of the actin core and reduction in actin in the foci. The decrease in myosin intensity is less pronounced in the same duration.

The length scale of the rips in the actin network determined whether the mode of contraction was homogeneous or coarse, with larger rips observed in the latter case. In both cases, actin intensity on the membrane reduced by 50% within a time range of 5±1.7 minutes after addition of myosin (Fig. S3A). The contraction events were myosin-specific and required a continued supply of ATP, as the process can be stalled at the bundling stage in the absence of ATP regeneration (Movie 4). Since myosin continues to bind actin in the absence of ATP, a catastrophic disintegration of the bundles was observed once new ATP was made available by the addition of an ATP regeneration mix.

The dense actin condensates formed during coarsening allowed a distinct observation of actin filament breakdown upon contraction. Representative images and kymograph of one of these foci illustrate the disassembly of actin filaments in the core after condensation (Fig. 2C a, b). Following the individual radial profiles of actin and myosin intensities in these foci at different time points, we observed that the condensation process was followed by a hollowing of the foci (Fig. 2D). The actin intensity at the core decreased drastically, while the myosin intensity showed a comparatively minor reduction (Fig. 2C b, 2D). This indicates that as contraction continues in the condensed actin foci, the actin filaments disintegrate rapidly under the stresses generated by the enriched myosin motors. A similar hollowing of actin condensates with myosin activity was described in solution experiments (Soares e Silva et al., 2011).

Individual events of filament or bundle fragmentation were difficult to observe in the dense network. However, as actin condensed into coarser foci, bundles of actin could be observed distinctly linking the foci. As contraction proceeded these bundles ruptured abruptly and recoiled, revealing the tensile stresses acting on them (Fig. S3B a, b). Myofilaments could be seen as distinct puncta along the bundles, presumably cross-linking the actin filaments (Fig. S3B c). This explicitly shows one mode of filament breakage occurring during contraction in the dendritic network.

### Redistribution in active networks results in a dynamic steady state of actin distribution

The stark contrast in actin distribution generated during coarsening also enabled us to observe phenomena which were not discernable during homogeneous contraction. Although coarsening initially generated a very heterogeneous actin distribution on the membrane, eventually the actin network resumed a seemingly uniform organization (Movie 5). In Figure 3A, we use the Coefficient of Variation (CV) of the intensity distribution of actin as an indicator of the heterogeneity in its physical distribution on the membrane. CV was calculated as the ratio of standard deviation to mean of the actin intensity distribution at each time point. The increase in CV coincided with the contraction process upon addition of myofilaments, peaking at the height of coarsening (Fig. 3A a-c). The CV then dropped as the actin was redistributed and finally results in an actin distribution nearly as homogeneous as before the addition of myosin (Fig. 3A d-f, graph). This observation suggests that large-scale heterogeneity in actin distribution is unstable in this dynamic minimal cortex. Under the combined influence of myosin-driven disassembly and actin polymerization, reorganization continues till a more homogeneous actin distribution is achieved. This distribution was similar to that observed when contraction proceeded homogeneously (Fig. 3A-graph).

**Figure 3.**
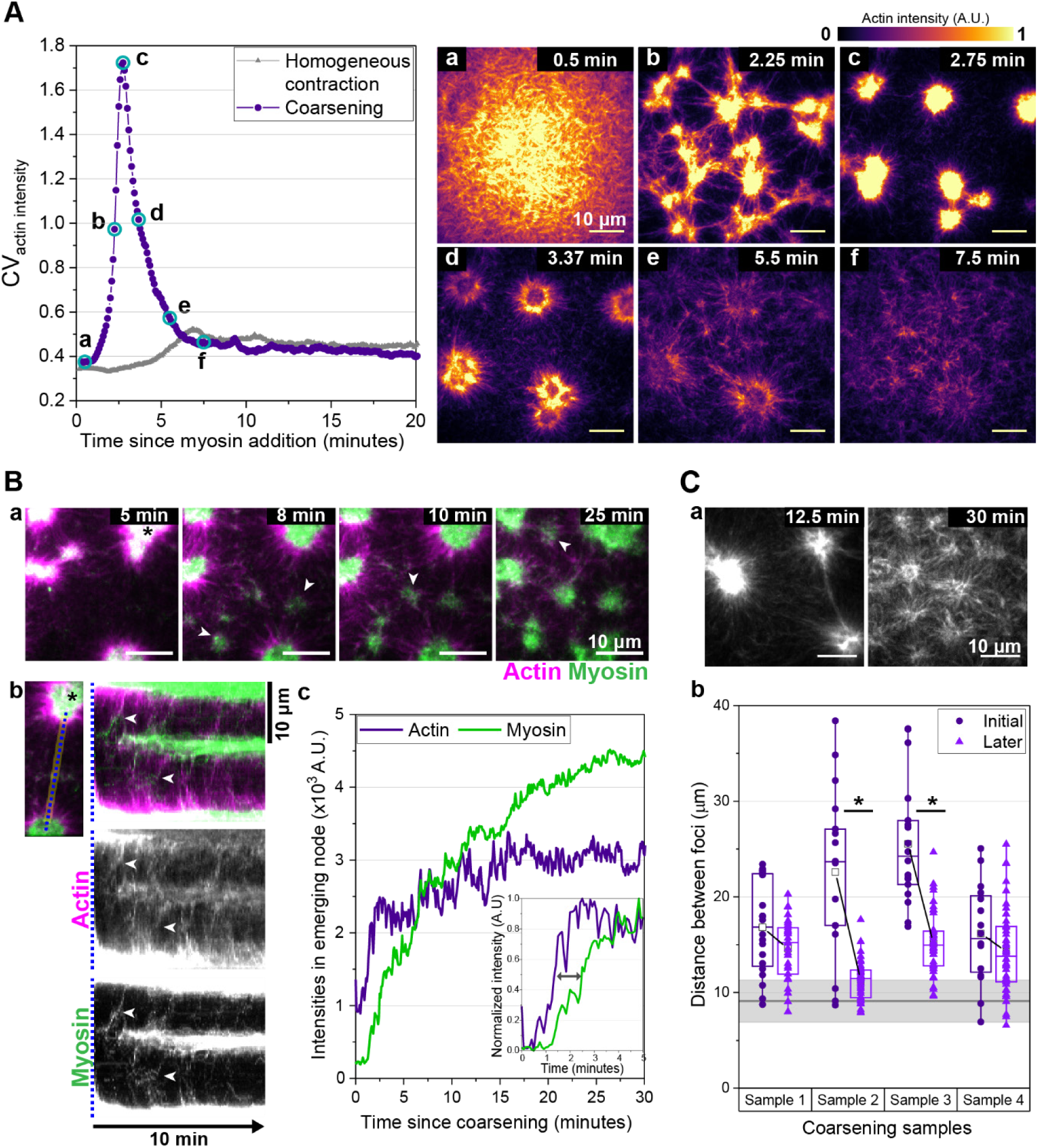
Redistribution in active networks restores homogeneous length scales. A) Heterogeneity of actin distribution during contraction is represented by the Coefficient of Variation (CV) of actin intensity in the image plotted over time. CV is calculated as the standard deviation divided by the mean for actin intensity distribution at each time frame. Images corresponding to the time points highlighted by cyan circles in the profile of coarsening are shown on the right (a-f). B) (a) The reappearance of actomyosin foci in regions where actin was removed during coarsening is highlighted by white arrows in selected time frames. (b) The kymograph plotted from a 10 pixel-thick line connecting two foci shows the emergence of new foci in between. White arrows indicate transfer of material between the foci. Actin and myosin are individually shown in grayscale in the lower frames. (c) The graph shows actin and myosin intensity changes in the newly emerging node measured in a circular region of 10-pixel diameter. The graph inset shows normalized intensities in the first 5 minutes as the new foci emerges. C) In samples showing coarsening, the distance between neighboring foci decreases over time, coming closer to the value of length scale of homogeneous contraction (gray line with the shaded area depicting mean±s.d.). Asterisk indicates samples were differences are significant after transition (p<0.05; Mann-Whitney test). Representative images of the transition of actin distribution in a sample are shown.

The restored uniformity in actin distribution can be attributed to two phenomena: the loss of actin from contraction foci and the recovery of actin filaments on the remaining regions of the membrane. These two processes occurred simultaneously, as can be illustrated by plotting the opposing trends in actin intensity in these two regions of the membrane (Fig. S4A). The loss of actin resulted from breakdown of condensed actin in the foci as described in the previous result (Fig. 2D) and also observable in Figure 3A (c-d). The concomitant reappearance of actin in other regions was accompanied by formation of new myosin foci in these parts (Fig. 3B a, white arrows, Movie 6). A kymograph of a stretch between two contraction foci revealed that both actin and myosin contribute to the newly emerging node (Fig. 3B b-c, data representative of 32 such nodes). Myosin intensity, however, increased with a slight delay after actin intensity in the new foci (Fig. 3B c, inset), suggesting that actin drives the reorganization. This delay was variable between nodes, ranging from 5 to 40 seconds. The source of the material for the new foci cannot be distinctly discerned, but both actin and myosin were exchanged with the neighboring foci (Movie 6, Fig. 3B b, white arrows). Myosin accumulated in the new foci with time, whereas actin intensity remained constant (Fig. 3B b, c). These observations indicate that myosin follows actin filaments and the redistribution of the former is limited as compared to the latter.

As can be expected, this reorganization resulted in a reduction in the distance between foci we previously measured as contraction length scale during coarsening (Fig. 3C a, b). The final values approached the length scale of homogeneous contraction (Fig. 3C), indicating that this actomyosin network has a favored length scale that is restored in spite of initial disturbances. The resultant actin distribution resembles a dynamic steady state with filament exchange between the foci observed even after an hour since addition of myosin (Movie 7, Fig. S4B).

Unlike myosin, VCA localization on the membrane was stable during contraction (Fig. S4C) and unaffected by coarsening (Fig. S4D), suggesting that VCA-driven actin polymerization could contribute to reappearance of actin during redistribution. Interestingly, VCA also enriched in the late-stage actomyosin foci formed on the membrane, possibly resulting in enhanced polymerization at these sites (Fig. S4D). However, while redistribution of actin resulted in local increase in actin intensity, mean actin intensity on the entire membrane did not increase after the initial reduction during contraction (Fig. S4E). Thus, a considerable proportion of the actin network is permanently lost from the membrane, coinciding with appearance of condensates in solution (Fig. S4F a), reminiscent of previous solution studies (Soares e Silva et al., 2011). The shape dynamics of these condensates indicate that myosin is active in them (Fig. S4F b), possibly trapping the actin filaments by cross-linking and recapture.

### New assembly of actin filaments reveals myosin-induced actin turnover in the minimal cortex

To investigate the role of myosin-driven fragmentation in actin filament turnover, we asked whether the network components released during myosin contraction can be reused for new polymerization. We reasoned that if excess monomers were washed off from the actin network before addition of myofilaments, any polymerization observed after myosin activity would largely be derived from the filament fragments and monomers generated by myosin-mediated breakdown of actin. To isolate the actin growth process, myosin-II activity was blocked using the inhibitor blebbistatin after the initial fragmentation and redistribution of actin had taken place. We found that the surface intensity of actin gradually increased after the inactivation of myosin (Fig. S5 a, b), suggesting that myosin breaks down the actin network to the level of monomers, which are then competent for new polymerization. In accordance with our hypothesis, novel polymerization was only observed if myosin had broken down the actin network, since no increase in actin intensity was observed if blebbistatin was added before actin network breakdown (Fig. S5 a, b). This indicates that breakdown by myosin facilitates new actin assembly by releasing monomers after network growth has saturated.

To follow the formation of new filaments in presence of myosin activity in greater detail, we designed a pulse-chase type experiment with two differently labeled actin monomers (Fig. 4A). The actin network was assembled as usual with 10% Alexa568-labeled actin (Label1). Excess monomers in solution were then removed by extensive washing. Along with myofilaments, additional actin monomers (1 μM) were added to the system, 10% of which were labeled with Alexa488 (Label2). Additional Arp2/3 complex was also provided so that the process of reassembly is not limited by its availability. We then observed whether the newly assembled actin on the membrane contained both labels.

**Figure 4.**
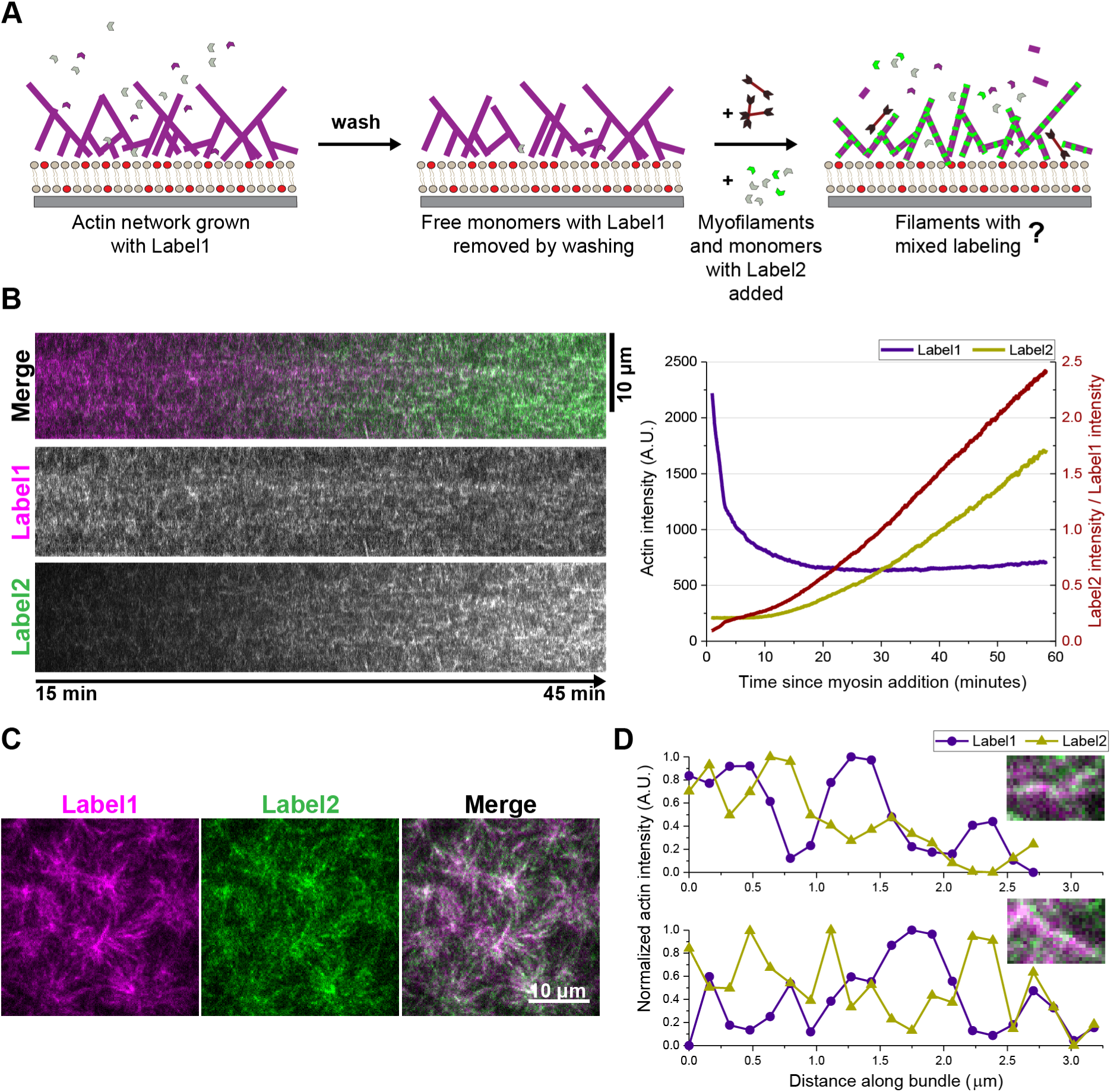
Actin undergoes turnover in the minimal dynamic cortex. A) Schematic of experiment design to follow the turnover of actin filaments observed on the membrane after myosin contraction. B) Left panel shows kymograph from an experiment of the type described in A. Label1 intensity shows a gradual reduction while the increase in Label2 shows a sharper increase. The right panel shows the corresponding graph showing intensity changes of the two labels in the actin network as well as the ratio of the two labels (dark red). C) Label2 appears as speckles on the actin structures observed at an intermediate time-point, indicating that the Label2 has an interspersed distribution on filaments with homogenous distribution of Label1. D) A 1-pixel wide line profile along two of the bundles in C showing normalized intensities of the two labels. The graphs appear to be anti-correlated, indicating that filaments of mixed labeling exist in these bundles.

We found that the proportion of Label2 in the membrane-associated actin increased over time after the initial breakdown of the Label1-network (Fig. 4B). This continuous increase confirmed that the actin filaments observed on the membrane were undergoing constant turnover in the presence of myosin activity. The steady increase in Label2 intensity on the membrane was accompanied by a very slight increase in Label1 intensity. The difference in the rates of change was probably due to the increase in total amount of actin on the membrane in this period and the total amount of Label2-actin in the system being higher.

In the intermediate stages of label mixing, filaments with uniform distribution of Label1 could be observed with speckled occurrence of Label2 (Fig. 4C). Since the actin on the membrane appeared bundled in the presence of myosin, individual filaments could not be distinguished. However, single pixel line profiles along two distinct bundles showed that the intensities of the two labels appear to be anti-correlated (Fig. 4D). Such filaments of mixed, interspersed labeling would only be observed if monomers of both Label1 and Label2 were available for polymerization, i.e. only if Label1 monomers were released from the pre-assembled network upon myosin contraction. Altogether, these observations suggest that the dynamic steady state actin distribution that emerges in this minimal cortex is comprised of actin filaments undergoing continuous turnover under the combined influence of new assembly of actin and myosin activity.

## DISCUSSION

In this study, we have reconstituted a minimal dynamic actin cortex to elucidate myosin-assisted turnover of actin filaments. We employed a TIRF microscopy approach to study the effects of myosin-II contraction on the reorganization of actively polymerizing branched actin networks on membranes. The key distinguishing feature of the minimal actin cortex described in this study is the coexistence of catalytic actin polymerization with the contractile activity of myosin. This enabled us to demonstrate myosin-II-stimulated actin turnover by observing the three essential processes of: 1) disassembly of pre-existing actin network, 2) redistribution of released network components, and 3) re-use of these components for new network assembly (Fig. 5A).

**Figure 5.**
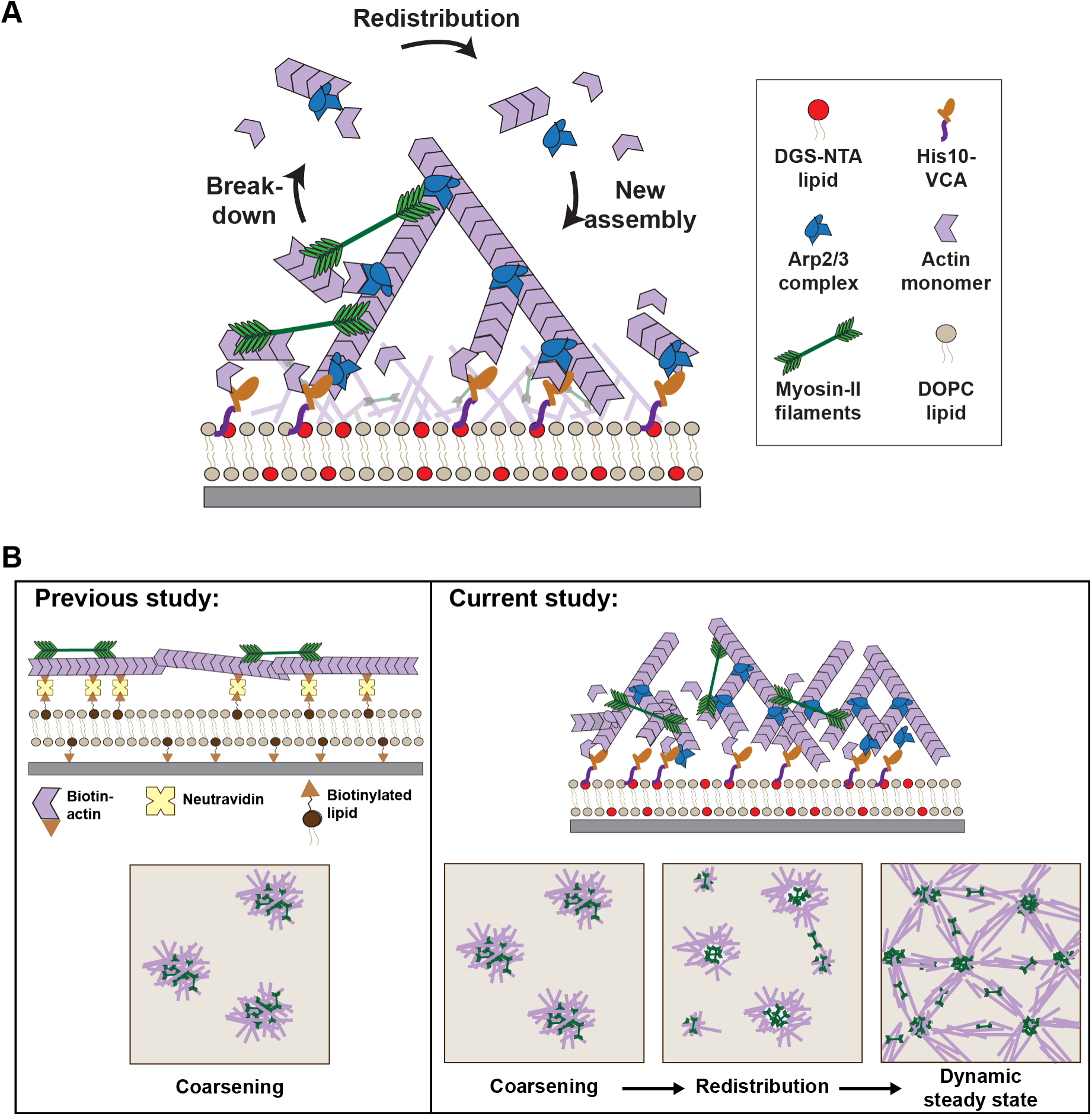
Myosin activity generates a dynamic steady state actin distribution with continuous turnover in a minimal actin cortex. A) Schematic summarizing the three key aspects of actin turnover in the minimal dynamic actomyosin cortex reconstituted in this study: 1) breakdown of membrane-bound branched actin networks by myosin, 2) redistribution of network components on the membrane, and 3) incorporation of these network components in the new actin assembly on the membrane. B) Schematic comparing the results of our study with a previous reconstitution of a minimal actin cortex, where pre-formed actin filaments were attached to an SLB via neutravidin-mediated bridging of biotinylated actin monomers and biotinylated lipids (Vogel et al. 2013). Our experimental design overcomes constraints on actin assembly and diffusion by using the VCA-Arp2/3 actin filament nucleation module that enables both fast actin polymerization as well as dynamic membrane attachment. Consequently, while myosin-II-driven reorganization of actin was limited to coarsening in that study, we observe a subsequent redistribution resulting in the emergence of a dynamic steady state actin distribution in our minimal dynamic actin cortex.

Altogether, our observations of myosin-driven reorganization of branched actin networks on membrane are well in line with previous studies, which also describe a distinct network coarsening upon myosin addition both in solution (Backouche et al., 2006, Kohler et al., 2011, Soares e Silva et al., 2011) or on membranes (Koster et al., 2016, Linsmeier et al., 2016, Murrell and Gardel, 2012, Smith et al., 2007, Stam et al., 2017, Vogel et al., 2013). However, beyond this condensation process, we further observed the subsequent de novo polymerization and redistribution of actin on the membrane (Fig. 5B), with only one of the solution studies previously reporting a similar phenomenon of an emergent dynamic steady state (Kohler et al., 2011). This finding highlights the dynamic nature of the minimal actomyosin cortex that we have reconstituted. In particular, the significance of including polymerization catalyzed by VCA is reinforced by the observation of less dynamic condensates in solution, as VCA is membrane-bound and not present in any significant amount in solution.

On a lipid bilayer, fragmentation of individual actin filaments by myofilaments, in isolation or within a dense network, has been demonstrated (Murrell and Gardel, 2012, Vogel et al., 2013). However, the breakdown and removal of membrane-bound actin after initial compaction was likely impaired by strong membrane adhesion of the biotinylated actin filaments (Vogel et al., 2013) or the use of crowding agents in solution (Murrell and Gardel, 2012). In contrast, in the minimal cortex described here, only the nucleation promoting factor (VCA) is firmly bound to the nickelated lipids in the membrane. The membrane association of the actin, resulting from the cumulative effect of multivalent interactions between the actin filaments and the VCA, is persistent but dynamic. This set-up is a reasonable approximation of cellular environments, where proteins linking actin networks to membrane undergo fast turnover (Fritzsche et al., 2014), effectively enabling a highly dynamic association by an alternate mechanism. Such reversible filament anchoring may be crucial for the myosin-driven disassembly of dendritic actin networks that we observe, as similar processes were reported when actin networks were grown from a related nucleation promoter, adhered to a glass surface (Reymann et al., 2012). Notably, we observed actin disassembly in the form of a distinct hollowing of actin condensates on the membrane, previously reported only in solution actomyosin aggregates (Soares e Silva et al., 2011). The solution study speculated that myosin coalesces in the centre of contraction foci when further translocation of myofilaments along the actin filaments is restricted by collision with other myofilaments. The polarized motion of myosin motors coupled with the capacity of actin filaments to reorient would normally result in the formation of a myosin-rich core with actin filaments extending outwards from it. The fragmentation of actin filaments under myosin-generated stress further allows a considerable fraction of the actin to disperse even if actin fragments remain associated with the myosin in the core, leading to an “actin shell” organization. However, in contrast with those observed in solution, these shell-like structures are transient in our minimal cortex, presumably because the polymerization machinery enables further redistribution of actin.

When coarsening was observed during contraction in our minimal cortex, some regions of the membrane were left largely devoid of actin. The resulting contrast in actin distribution enabled direct observation of the redistribution of network components on the membrane. Synergistic relocation of actin and myosin, facilitated by actin polymerization, led to a more homogeneous distribution of contraction foci. The active network then continued to show long-term flows of actin filaments over this preferred length scale, resulting in the emergence of a dynamic steady state of actin distribution, previously described only in solution clusters (Kohler et al., 2011). Such fluidization of membrane-associated actin networks by motor activity may aid the fast shape dynamics of the eukaryotic actin cortex.

The transition of length scale of actomyosin dynamics that results from redistribution provides a compelling avenue for further investigation. While we offer some preliminary analyses of the factors that affect the length scale of actomyosin contraction, further study is needed to determine why our cortical active network shows bimodal length scales. We observed a dependence on net membrane association of the actin network, branching densities and possibly filament length and active motor concentration, consistent with previous observations during contraction of disordered actin networks on membrane (Murrell and Gardel, 2012). These are all factors that are subject to extensive biochemical regulation in cell cortices and could, therefore, contribute to the coordination of actomyosin activity over different length scales in a cell.

Finally, we elucidate the contribution of myosin activity in actin filament turnover in this reconstituted cortex with the direct observation of the phenomenon. The notion of myosin enhancing actin turnover rates has been suggested in various cellular contexts (Guha et al., 2005, Medeiros et al., 2006, Murthy and Wadsworth, 2005, Yang et al., 2012, Wilson et al., 2010), but the inhibition of myosin activity is a fairly complex perturbation due to the many roles of myosin-II motors in the cytoskeleton. By disassembling actin networks, myosin can amplify local cortical remodeling by providing fast access to a pool of network components for new assembly. Since actin organization is to be coordinated over micron-scale distances in the cell from a limited pool of components, this suggests an interesting mechanism by which myosin activity could support faster dynamics on a shorter scale. Furthermore, the negative feedback on contractility due to disassembly could form the basis of the pulsatory dynamics observed during certain developmental processes (Nishikawa et al., 2017, Martin et al., 2009).

In this simplified cortex, we selected only the minimal components required for dendritic network assembly and contraction to gain a clearer understanding of the role of each participating component. Altogether, this study demonstrates that myosin activity, when counterbalanced by actin polymerization, can induce actin turnover in a minimal cortex and result in the formation of an active network in dynamic steady state on the membrane. An interesting avenue for more detailed investigation is to further dissect the actomyosin organization in the emergent distribution and determine the influence of other structural components of cortical networks. A detailed comparison with non-muscle myosin-II in a similar set-up will also enhance our understanding of a cellular setting. Importantly, our experiments suggest that the reconstituted dynamic actin cortex described in this study is readily amenable to increased complexity and controlled manipulations. It, therefore, holds great potential for providing further insights into the functioning of its biological inspiration.

## MATERIALS AND METHODS

### Proteins and other reagents

VCA domain (amino acids 400-501) of murine N-Wasp (Uniprot ID Q91YD9) was cloned into a modified pCoofy vector with 10x Histidine tag. The protein was then purified by a combination of affinity and anion exchange chromatography. In brief, the protein was expressed in a 1 L culture of Rosetta T1 strain of BL21 cells by inducing with 0.5 mM IPTG and growing the cells for 16 hours at 16°C. For purification, cells were lysed by sonication. The cell lysate was purified with the His tag on a HisTrap HP column (GE Healthcare, Chicago, IL, USA) via a chromatography system. The protein containing fractions were combined and diluted 10 times for ion exchange chromatography on a MonoQ column (GE Healthcare, Chicago, IL, USA). The selected protein fractions were buffer exchanged into phosphate buffer saline supplemented with 10% Glycerol. All buffers included 1 mM TCEP to prevent protein dimerization.

VCA was labeled with Atto488-maleimide (28562) from Sigma-Aldrich Corporation, St. Louis, MI, USA. The dye was used in 20-fold excess of the protein (by mass) and the mix was incubated overnight in a cold room with gentle rotation. Buffer exchange was done on a small volume Sephadex G-25 column. This was followed by dialysis for complete removal of the dye. The final storage buffer of the protein was 50 mM Tris-HCl pH 7.5, 150 mM NaCl, 10% Glycerol and 1 mM TCEP. Labeling efficiency was quantified by spectrophotometric analysis.

Purification of myosin-II purification from rabbit muscle, as well as labeling with Alexa Fluor 488, was carried out as described previously (Vogel et al., 2013).

Unlabeled actin monomers from rabbit skeletal muscle (AKL99) and Arp2/3 complex from porcine brain (RP01P) were purchased from Cytoskeleton, Inc., Denver, CO, USA. Actin monomers conjugated with Alexa Fluor 568 (A12374) and Alexa Fluor 488 (A12373) were acquired from ThermoFischer Scientific, Waltham, MA, USA. Capping protein CapZ (non-muscle, human recombinant) was purchased from HYPERMOL EK, Germany.

Blebbistatin (B0560), Phosphocreatinine (P7936), Creatine Phosphokinase (C3755), Pyranose oxidase (P4234) and Catalase (C40) were purchased from Sigma-Aldrich Corporation, St. Louis, MI, USA.

### Preparation of lipid bilayer

DOPC and DGS-NTA lipids were purchased from Avanti Polar Lipids, Inc., Alabaster, AL, USA. Small unilamellar vesicles with desired proportion of DOPC:DGS-NTA lipids (with 0.005% Atto655-DOPE) were prepared by rehydrating a lipid film in SLB buffer (50 mM Tris-HCl pH 7.5, 150 mM KCl) followed by sonication. The composition was 99 mol% DOPC and 1 mol% DGS-NTA in all experiments, unless otherwise mentioned. High precision coverglass (no. 1.5, Marienfeld-Superior, Lauda-Königshofen, Germany) was treated with Piranha solution (3:1 H_2_SO_4_:H_2_O_2_, overnight), rinsed with copious amounts of milliQ water, dried and subsequently briefly treated with O_2_ plasma (Femto plasma cleaner, Diener-plasma, Ebhausen, Germany). Reaction chambers were generated on these coverglasses by press-to-seal silicone isolators (GBL664206; Grace Bio-labs, Bend, OR, USA). 0.1 mg/ml of vesicles were deposited in the silicone chambers with frequent pipetting, followed by 15 minutes of incubation. The bilayers were gently washed first with SLB buffer and then with VCA buffer (10 mM Tris-HCl pH 7.5, 50 mM NaCl, 1 mM DTT) and stored in this buffer till sample preparation.

### Sample preparation

A lipid bilayer was flushed again with VCA buffer and then incubated with 200 nM 10xHis-VCA in the same buffer. For experiments where labeled VCA was required, Atto488-labeled VCA was included in a 1 in 10 ratio with unlabeled VCA. A premix containing actin monomers (10 % Alexa568-labeled), Arp2/3 complex and ATP regeneration system (Phosphocreatine and Creatine Phosphokinase) was made at the same time in G-buffer (10 mM Tris-HCl pH 8.0, 0.2 mM CaCl_2_, 0.2 mM ATP, 1 mM DTT). After 20 minutes, the excess VCA was washed off and actin pre-mix was added. An image was taken for the reference intensity value for normalization before triggering polymerization by adding the 5x G-to-F buffer (50 mM Tris-HCl pH 7.5, 250 mM KCl, 10 mM MgCl2, 1 mM DTT with variable ATP). A typical sample finally contained 1 μM actin monomers and 10 nM Arp2/3, unless otherwise mentioned in the figures. Additionally, the sample also included the ATP regeneration system (20 mM Phosphocreatinine, 53 U/ml Creatine Phosphokinase), and an oxygen scavenger system (37 U/ml Pyranose Oxidase, 90 U/ml Catalase and 0.8 % Glucose). The final buffer conditions were 10 mM Tris-HCl pH 7.5, 50 mM KCl, 1mM MgCl2, 1mM DTT with ATP adjusted to 0.1 mM. The sample volumes were 40 ml.

Myofilaments were pre-assembled by making a 4x stock of the myosin in a buffer of 10 mM Tris pH 7.5, 50 mM KCl, 2 mM MgCl_2_, 1 mM DTT and incubating for 10 minutes before adding to the actin network by replacing ¼ of the sample volume. Where washing was required before addition of myosin, a 4-fold replacement of sample volume was carried out with gentle mixing.

In all cases, evaporation was minimized by storing the samples in closed boxes with wet tissue till they were used. For putting on the microscope, a hydrophobic pen was used to draw a "moat" around the silicone well and this was filled with water. A small plastic lid was used to cover the sample, including the moat, during imaging.

Variations from this basic protocol have been mentioned in the text, where relevant. Blebbistatin was dissolved in DMSO and used at a final concentration of 20 μM.

For experiments in which a second label of actin was needed, Alexa488-labeled actin monomers were used. For pulse-chase experiments in Figure 4, Alexa488-labeled monomers were mixed with unlabeled actin monomers at a 1 in 10 ratio. 1 μM of this 10 % labeled actin was added to the sample, along with 10 nM Arp2/3, as described in the text.

### Image acquisition

Fluorescence images were recorded on a home-built objective-type TIRF microscope (Mucksch et al., 2018), constructed around a Nikon Ti-S microscope body with oil immersion objective (Nikon SR Apo TIRF, 100x, NA 1.49, Nikon GmbH, Düsseldorf, Germany). In brief, the excitation laser lines (490 nm (Cobolt Calypso, 50 mW nominal), 561 nm (Cobolt Jive, 50 mW nominal) and 640 nm (Cobolt 06-MLD, 140 mW nominal)) were attenuated with an acusto-optical tunable filter (Gooch & Housego TF-525-250) and spatially filtered by a polarization-maintaining single-mode fiber (kineFLEX-P-3-S-405.640-0.7-FCS-P0 and kineMATIX, Qioptiq, Hamble, UK). After the fiber, the excitation beam was collimated, extended 3-fold by a telescope of achromatic doublets and moved off the optical axis by a mirror and a focusing lens, both on a piezo-controlled translation stage. This way, the excitation beam was focused off-axis on the back focal plane of the objective. Excitation and emission light were separated using a four-color notch beam splitter (zt405/488/561/640rpc flat, AHF Analysentechnik, Tübingen, Germany). For single-color detection, the fluorescence light was going through a 4f lens system, including spectral filtering (525/50, 593/46, 705/100) and detected on an electron-multiplying charge-coupled device (EMCCD) camera (iXon Ultra 897, Andor Technologies, Belfast, UK). For dual-color schemes, the image was laterally clipped in a conjugated focal plane. The clipped images were spectrally separated (T555LPXR, Chroma), bandpass filtered (525/50, 593/46) and focused side-by-side on the same EMCCD. To minimize phototoxicity and –bleaching, the AOTF transmission was synchronized with the EMCCD acquisition using a controller card (PCle-6323, National Instruments) with a customized LabView software (National Instruments, Austin, USA). Imaging interval was either 2 or 5 seconds, with exposure times of 50 or 100 msec. A custom-built focus stabilization eliminated drift of the focus position, based on a feedback loop, which monitored the position of an infrared laser beam, back-reflected from the sample interface.

For imaging in solution, as shown in Figure S12, a Yokogawa scan head CSU10-X1 spinning disk system set up on a Nikon Eclipse Ti inverted microscope body was used. Detection was done with an EMCCD camera (iXon Ultra 897, Andor Technologies, Belfast, UK) and a 3i solid state diode laser stack with 488 nm, 561 nm and 640 nm laser lines (3i, Denver, Colorado, USA) was used for excitation. Imaging was done with a UPLanSApo 60x/1.20 Water UIS2 objective (Olympus, Japan).

### Image processing and analysis

ImageJ (Rueden et al., 2017) was used for image processing and analysis. Rolling ball background subtraction was used to highlight features in Figure 2A. Rolling ball radii were 100 pixel for actin and 20 pixel for myosin.

Customized macros were made for most intensity and length scale analyses. For intensity measurement, the macro included a module to specify a square region (edge length 200 pixel) of the sample and then measure mean intensity and standard deviation for each frame. For calculation of Coefficient of Variance in Figure 3, the ratio between the mean intensity and the standard deviation was calculated for each time frame. The macro for length scale analysis combined the use of multipoint selection and the mathematical transformation of the pixel positions to lengths in microns. Additional plugins used were Radial_Profile_Angle_Ext and Kymo_wide_reslice (available for download from https://imagej.nih.gov). The radial intensity profiles in Figure 2D were made by using the Radial_Profile_Angle_Ext plugin, which integrates the intensities at each point along the radius of a circular selection. The profile was mirrored to plot the average intensity along the diameter for aiding visualization of the node organization. For Kymo_wide_reslice, the pixel thickness of the selected line is specified with corresponding data in the figure legend.

The software FFmpeg was used for making movies from time series data (available for download from http://ffmpeg.org/).

Statistical analysis and plotting of graphs was done in Origin. Fitting of curves was also done in Origin using the in-built model “ExpGrow1”. Fitting excluded the initial 3 minutes for achieving a better fit. For significance testing in length scale measurements, values were pooled across samples for each condition. The nonparametric Mann-Whitney test was used to compare two populations at a 0.05 significance level.

## ACKNOWLEDGMENTS

The authors would like to thank Prof. Klemens Rottner for sharing plasmids and Sigrid Bauer, Dr. Katharina Nakel, Michaela Schaper and Kerstin Andersson, for technical assistance. This work is partially supported by MaxSynBio Consortium, which is jointly funded by the Federal Ministry of Education and Research of Germany and the Max Planck Society. KAG has received funding from the European Union’s Horizon 2020 research and innovation programme under the Marie Skłodowska-Curie grant agreement No. 703132. JM is grateful for financial support from the excellence cluster Nanosystems Initiative Munich. S, JM and PB acknowledge support from the International Max Planck Research School for Molecular and Cellular Life Sciences (IMPRS-LS). JM and PB acknowledge support from the Center for NanoScience (CeNS).

## LIST OF ABBREVIATIONS

Arp2/3: Actin-related protein 2/3
ATP: Adenosine Triphosphate
CV: Coefficient of Variation
DGS-NTA: 1,2-Dioleoyl-sn-glycero-3-phosphoethanolamine
DOPC: 1,2-Dioleoyl-sn-glycero-3-phosphocholine
DTT: Dithiothreitol
IPTG: Isopropyl β-D-1-thiogalactopyranoside
NPF: Nucleation Promoting Factor
s.d.: Standard deviation
SLB: Supported Lipid Bilayer
TCEP: tris(2-carboxyethyl)phosphine
TIRF: Total Internal Reflection Fluorescence
VCA: Verpolin homology, cofilin, and acidic domain

